# Holistic in silico developability assessment of novel classes of small proteins using publicly available sequence-based predictors

**DOI:** 10.1101/2023.11.10.566515

**Authors:** Daniel A. M. Pais, Jan-Peter A. Mayer, Karin Felderer, Maria B. Batalha, Timo Eichner, Sofia T. Santos, Raman Kumar, Sandra D. Silva, Hitto Kaufmann

**Affiliations:** Valgenesis Portugal, Lda, Avenida António Augusto Aguiar, 108, 4^th^ floor 1050-205, Lisboa, Portugal; Pieris Pharmaceuticals GmbH, Zeppelinstr. 3, 85399 Hallbergmoos, Germany

**Keywords:** Developability, in silico prediction, machine learning, early-stage drug development, small protein therapeutics

## Abstract

The development of novel therapeutic proteins is a lengthy and costly process, with an average attrition rate of 91% (Thomas et al., 2021). To increase the probability of success and ensure robust drug supply beyond approval, it is essential to assess the developability profile of new potential drug candidates as early and broadly as possible in development (Jain et al., 2023). Predicting these properties *in silico* is expected to be the next leap in innovation as it would enable significantly reduced development timelines combined with broader screens at lower costs. However, developing predictive algorithms typically requires substantial datasets generated under very defined conditions, a limiting factor especially for new classes of therapeutic proteins that hold immense clinical promise.

Here we describe a strategy for assessing the developability of a novel class of small therapeutic Anticalin® proteins using machine learning in conjunction with a knowledge-driven approach. The knowledge-driven approach considers developability attributes such as aggregation propensity, charge variants, immunogenicity, specificity, thermal stability, hydrophobicity, and potential post-translational modifications, to calculate a holistic developability score. Based on sequence-derived descriptors as input parameters we established novel statistical models designed to predict the developability scores for Anticalin proteins. The best models yielded low root mean square errors across the entire dataset and were further validated by removing input data from individual screening campaigns and predicting developability scores for those drug candidates.

The adoption of the described workflow will enable significantly streamlined preclinical development of Anticalin drug candidates and could potentially be applied to other therapeutic protein scaffolds.

## Introduction

Therapeutic proteins, a class of therapeutics produced in living cells, revolutionized modern medicine (Kesik-Brodacka, 2018). They exhibit high specificity and efficacy, have fewer side effects, and their high activity, long half-life, and low immunogenicity allow the treatment of a variety of severe diseases (Kesik-Brodacka, 2018; O’Flaherty et al., 2020), some of which were previously considered untreatable.

The dominant class of biologics for many years has been IgG-based therapeutic antibodies. Their ability to recognize and bind to specific antigens with high affinity enabled numerous clinical applications, from immune checkpoint inhibitors in cancer therapy to immune modulators in autoimmune diseases (Walsh, 2018). However, monoclonal antibodies have several limitations in clinical applications: their large size of around 150 kDa limits tumor penetration (Shah & Betts, 2013; Thurber et al., 2008), and their planar paratopes make it challenging to engineer binders against grooves or catalytic sites of enzymes (Skerra, 2000). These factors led to the development of several new therapeutic modalities such as single chain variable fragments and nanobodies (Kesik-Brodacka, 2018), and gave rise to novel classes of therapeutics based on engineered protein scaffolds (Gebauer & Skerra, 2020). Among those next generation-therapeutics are Anticalin proteins, a class of clinical stage biopharmaceuticals that enhance the range of drug formats and applications, and thereby address several limitations of monoclonal antibodies.

Anticalin-based therapeutic proteins are derived from lipocalin proteins, a family of extracellular proteins found in all kingdoms of life, except for Archaea (Ganfornina et al., 2022). Despite low sequence homology, lipocalins share a common architecture, characterized by an eight-stranded antiparallel beta-barrel fold, which forms an internal cavity (Flower et al., 1993) flanked by four flexible loops. This topology gives rise to many biological functions, including the storage, transport, and sequestration of ligands, which can vary from small molecules to large proteins. These structural features of the lipocalin fold offer important advantages for the development of therapeutics, such as format flexibility, stability, and potency, leading to the establishment of a modular and potent therapeutics’ platform (Morales-Kastresana et al., 2022). Recently, the development of antibody-Anticalin fusion proteins (Mabcalin™) and their application in treating various forms of cancer has been described (Hinner et al., 2019; Peper-Gabriel et al., 2022). Several Mabcalin proteins have been designed to combine T-cell stimulation mediated by 4-1BB agonism with various tumor targets or checkpoint blockade via PD-L1. Their successful advancement to ongoing clinical studies has facilitated the development of an efficient and robust manufacturing platform (Wachter et al., 2023).

The feasibility of turning a promising biological drug candidate into a viable, market-ready therapeutic is influenced by numerous factors. To deliver a high-quality drug to patients, a robust manufacturing process and stable patient-friendly dosage forms are crucial. Therefore, important drug developability properties such as production yield, aggregation propensity, charge variants, thermal stability, immunogenicity, to name a few (Bailly et al., 2020; Hartmann & Kocher, 2015; Narayanan, Dingfelder, Butté, et al., 2021; Xu et al., 2019) need to be assessed during the early stages of the drug development process and need to match the defined target product profile (TPP).

A robust manufacturing process seamlessly integrates risk assessment, risk mitigation strategies, and knowledge management together with a thorough analytical evaluation of each candidate. This holistic approach enhances the probability of successful product development by ensuring that potential challenges are identified early, thereby streamlining the path from concept to market-ready product (Bailly et al., 2020; Hartmann & Kocher, 2015; Jarasch et al., 2015). However, such strategy can only be successfully pursued if the desired characteristics of the developability profile of an early drug candidate are clearly defined and predictive for later stages of the drug life cycle.

An increasing body of literature describes experimental strategies to assess the overall developability profile of biologic drug candidates (Bailly et al., 2020; Chen et al., 2020; Goyon et al., 2017; Hartmann & Kocher, 2015; Jarasch et al., 2015; Raybould et al., 2019; Xu et al., 2019). A typical approach to drug development would be based on very limited analytical assessments using high throughput assays at early discovery stages with many hundreds of candidates. The analytics would then be broadened during lead optimizations focusing on a handful of leads (Jarasch et al., 2015; Khetan et al., 2022; Narayanan, Dingfelder, Butté, et al., 2021).

Recently, several teams have developed predictive tools for different attributes of protein drug candidates, and machine learning algorithms combined with *in silico* methods have proven to be a powerful approach (Bailly et al., 2020; Chen et al., 2020; Hebditch & Warwicker, 2019; Khetan et al., 2022; Lauer et al., 2012; Raybould et al., 2019; Tiwari et al., 2022). Machine learning is a sub-field of artificial intelligence that is focused on the development and usage of mathematical algorithms and statistical models that can learn from data and make predictions or decisions without being programmed explicitly. By applying such algorithms to a dataset containing multidimensional inputs and outputs, the algorithms can capture the patterns present in the input information that are correlated to the outputs, allowing to develop predictive models for the desired output (Narayanan, Dingfelder, Butté, et al., 2021; P. Schneider et al., 2020). Still, the use of machine learning tools for early candidate screening is not a one-size-fits-all solution, but rather an “augmented intelligence” endeavor with an emphasis on process knowledge (P. Schneider et al., 2020).

The use of machine learning and artificial intelligence in the advancement and development of biologics has been reviewed by various authors (Narayanan, Dingfelder, Butté, et al., 2021; Nikita et al., 2022; P. Schneider et al., 2020). Described applications include antibody design (Akbar et al., 2022), protein engineering (Yang et al., 2019), formulation development (Narayanan, Dingfelder, Condado Morales, et al., 2021), prediction of protein function including antigen-antibody binding from structure based models or primary amino acid sequence (Bileschi et al., 2022; C. Schneider et al., 2022), and prediction of biophysical properties such as melting temperature or methionine oxidation from antibody amino acid composition or protein structure (Gentiluomo et al., 2019; Sankar et al., 2018).

Most of the work to date on predicting developability attributes such as aggregation propensity or more general developability profiles focused on sequences based on the IgG subclass of proteins (Lauer et al., 2012; Obrezanova et al., 2015; Raybould et al., 2019). Developing such algorithms is based on the existence of large publicly available datasets linking analytical characterization (as exemplified in (Jain et al., 2017), to antibody sequence, structure, and antigen binding information (compiled in (Khetan et al., 2022)).

However, while classical IgG-based therapeutics gave rise to the majority of approved biopharmaceutics on the market, the emerging R&D pipelines are fueled by innovative formats such as bispecifics and small protein scaffolds (Löfblom et al., 2011; Mehta & Cochran, 2017). Here, much less experimental data is available to feed predictive models.

In this study we set out to define a comprehensive knowledge- and data-driven developability score for an innovative class of small Anticalin proteins following Quality by Design principles (ICH, 2009). The developability score approach devised herein uses both experimental data and *in silico*-predicted developability attributes, summarizing the multivariate information from analytical variables deemed critical for developability into one variable only (i.e., the developability score). Based on this methodology and using the most comprehensive experimental dataset available for this class of molecules, we developed *de novo* predictive models aiming to predict the developability scores based on amino acid sequence attributes. These models only depend on *in silico* parameters derived from the amino acid sequence, thus lowering the costs associated with analytical characterization of new drug candidates by reducing the number of candidates to be screened and analytical measurements needed.

## Materials and methods

### Dataset

The dataset used consists of 136 Anticalin protein candidates based on the neutrophil-gelatinase associated lipocalin (NGAL) scaffold and derived from screening campaigns against 5 different targets. Anticalin proteins were produced in *E. coli* with a C-terminal Strep-tag II and purified via Strep-Tactin XT affinity chromatography followed by preparative size-exclusion chromatography and Mustang E filtration. The candidates were analyzed via analytical size-exclusion high-performance liquid chromatography (SEC-HPLC), dynamic and static light scattering (DLS/SLS), capillary isoelectric focusing (cIEF), Heparin high-performance liquid chromatography (Heparin-HPLC), hydrophobic interaction high-performance liquid chromatography (HIC-HPLC), thermal scanning fluorimetry (TSF) and off-target binding to a comprehensive panel of proteins using a Luminex® multiplex assay (off-target Luminex) as well as off-target binding to target-negative cells (off-target FACS).

### *In silico* calculations

#### Potential post-translational modifications

For each Anticalin protein construct, potential post-translational modifications were counted based on the protein amino acid sequence using an *in house* script. The motifs considered for PTM counts are detailed on Supplementary Table 1.

#### Immunogenicity

Immunogenicity was calculated based on the protein amino acid sequence using the Epibase® platform.

#### Homology-based descriptors

Homology model-based descriptors were calculated using the Molecular Operating Environment (MOE) software (MOE, 2022).

#### Amino acid sequence-based descriptors

Protein sequence-based descriptors were calculated using the publicly available library protr (R package version: 1.6-2)(Xiao et al., 2015). For the three autocorrelation descriptors (Normalized Moreau-Broto, Moran and Geary) and the quasi-sequence order descriptors, a maximum lag distance of 30 amino acids was used. For the pseudo-amino acid composition descriptors, the lambda value of 30 was chosen. For further detail on the calculation of the protr descriptors and the choice of lambda and maximum lag distance, the reader is referred to (Xiao, 2015; Xiao et al., 2015).

#### Meta-variables

Two additional meta-variables were calculated, based on the difference between the measured value of biophysical measurements and its difference to the theoretical sequence-based values. This was performed for isoelectric point and molecular weight variables.

### Criticality assessment

The developability attributes were defined based on our internal knowledge of Anticalin protein development and antibody literature. Ultimately, seven distinct developability attributes were deemed critical for the development of Anticalin proteins: aggregation propensity, charge variants, immunogenicity, specificity, thermal stability, hydrophobicity, and potential post-translational modifications.

For the critical variables, a Quality Risk Management-based criticality assessment was conducted to evaluate the impact of each variable on four key product developability areas: manufacturability, stability, safety, and efficacy. Uncertainty regarding the impact was also factored into the criticality assessment. According to the combined uncertainty/impact score, variables were attributed a criticality score, with variables scoring higher than a predefined threshold being considered critical. On top of the criticality assessment, previous knowledge on the variables was also considered to define a variable as critical or non-critical. Each critical variable was assigned to one developability attribute, according to the potential of that variable to work as a surrogate for that attribute.

### Data preprocessing and scoring approach

Most modeling algorithms cannot deal with missing data. Herein, two missing data imputation algorithms were tested: MICE (multiple imputation by chain equation) and IPCA (Iterative PCA). The computations by both approaches were very similar. However, for some variables IPCA imputation yielded negative values in a positive only variable, and consequently missing data was imputed with the MICE algorithm (Azur et al., 2011), taking advantage of the algorithm’s flexibility. Imputation was only applied to variables with less than 20% of missing data. Additionally, many machine learning algorithms, including partial least squares (PLS), benefit from data transformation (James et al., 2021). Several transformation functions were investigated to transform strongly non-normally distributed variables. In the end, two variables were transformed using a cubic or negative reciprocal transformations, resulting in a reasonably reduced score for values considered as unfavorable as well as in a variable distribution closer to the normal distribution. As a final step in exploratory data analysis, a multivariate outlier removal step was performed, considering the construct’s distribution in the principal component analysis (PCA) score plot and the Hotelling’s T^2^ and Q-Residuals metrics (Thennadil et al., 2018).

To calculate the developability score for each construct, the analytical values for each original variable or transformed variable were normalized into a score, ranging from 0 to 10, with 10 being the optimal value for that variable. The optimal and pessimal values were defined as either the minimum or maximum value for that variable observed in the dataset, depending on whether an “optimal construct” would have a high or a low value in that variable, according to the defined developability target profile (Figure 1, “Max/Min” column). Afterwards, the construct’s developability score was calculated as the average of all normalized values for the 17 critical variables.

**Figure 1.**
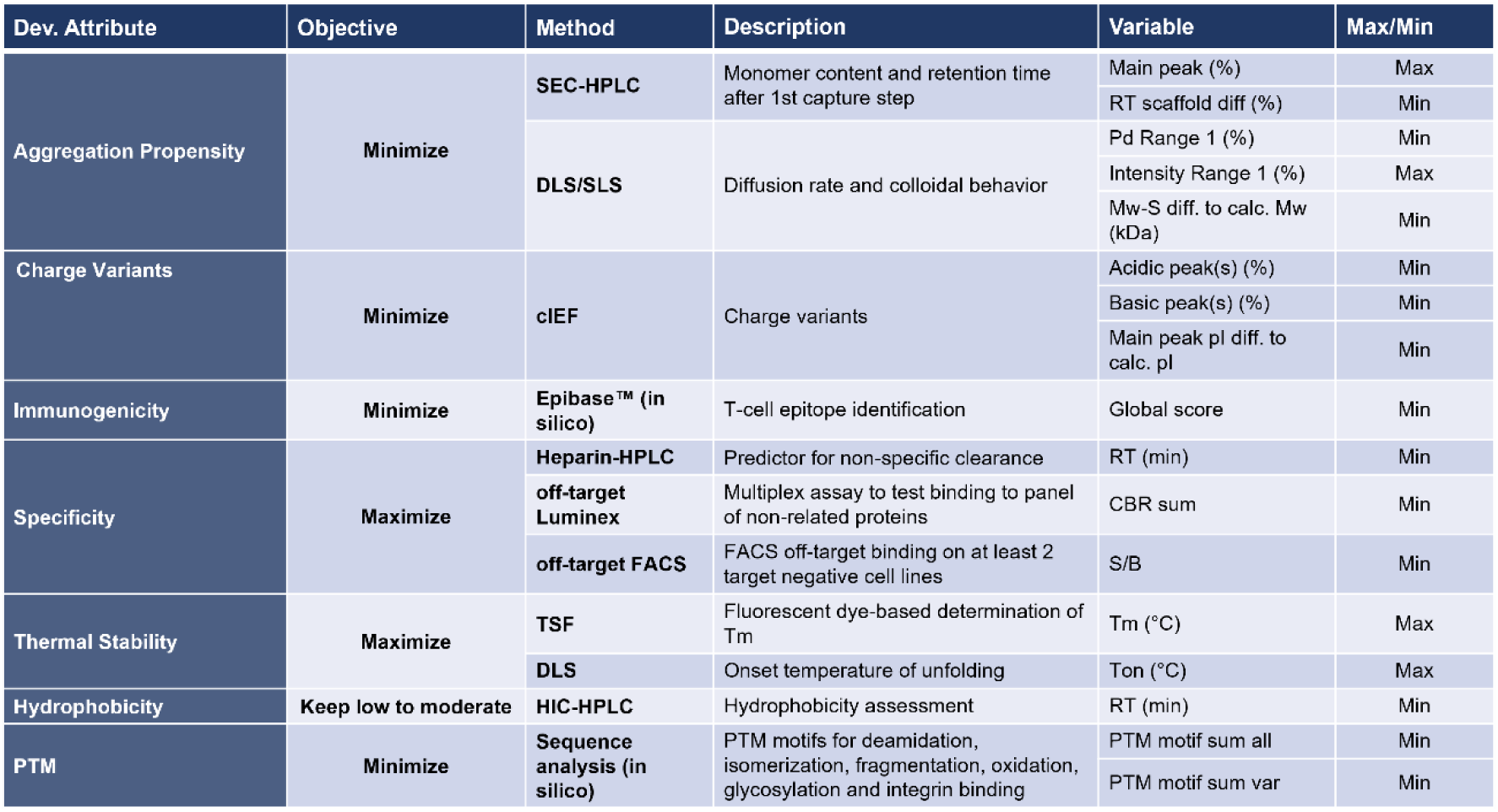
Developability attributes, critical variables, and objectives for the developability scoring approach. For each developability attribute, an objective (maximize or minimize) and variables that can serve as surrogate to measure that developability attribute are defined. To perform variable normalization as described in materials and methods, for each variable the “max/min” target is defined.

### Modeling strategy

All calculations and modeling activities were performed using R (version 4.0.3) and RStudio (version 2022.12.0 Build 353). PCA models were computed using the mdatools package (version: 0.12.0) (Kucheryavskiy, 2020).

*In silico* descriptors were obtained from the publicly available library protr (Xiao et al., 2015) and by using the Molecular Operating Environment (MOE) software (MOE, 2022). Protr is an open-source R toolkit for protein sequence-derived structural and physicochemical descriptor computation. It generates numerical representation schemes of protein sequences based on the AAindex database (Kawashima et al., 2007) which can be generally divided in various groups and families of descriptors, e.g., 240 normalized Moreau-Broto descriptors. MOE descriptors were determined by generating a template-based 3D homology model for each construct and subsequently calculating features like e.g., the total area and area of the largest hydrophobic, positively and negatively charged patches.

To correlate the *in silico* descriptors with the calculated developability scores, both PLS and support vector machine regression (SVM) were tested using the caret package (version: 6.0-93) (Kuhn, 2008). The dataset was divided into a training (75%) and a validation (25%) set. To prevent target overrepresentation in either the training or validation set, the division was performed according to the Anticalin target representation within the dataset (e.g., if a given target was 40% of the whole dataset, that target was kept approximately at 40% in the training and validation set as well). The number of PLS latent variables and the SVM parameters (polynomial degree, regularization parameter “C” and scaling parameter) were determined using cross-validation in the training set, with a 10-fold cross-validation and 3 repeats. For PLS, the latent variables tested ranged from 1 to the minimum between 50, number of variables or number of constructs of the dataset. For SVM, the default caret parameters were used (a grid search between the 3 SVM hyperparameters). The model with the lowest cross-validation root mean squared error (RMSE) was then used for predicting the validation set.

To optimize the number of descriptors, recursive feature elimination (RFE) was performed with resampling using the caret package. RFE is an iterative process in which a model is built using all variables, the variable importance is calculated and then the less informative variables are removed; this process is then iteratively repeated until a predefined number of variables is reached. It assures that only uninformative variables are removed, and in the end only the informative variables are kept.

RFE was performed for a pre-defined sequence of different number of variables, ranging from 10 up to the maximum number of predictor variables existent in the dataset, in increments of 100 variables. The RMSE of cross-validation was calculated for each model obtained. A moving average and standard deviation were computed for the last 3 models. The final number of variables was then selected based on the value of 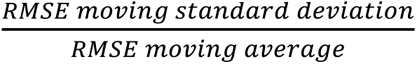, considering a maximum threshold of 2% for the latter ratio.

The chosen model was then trained with the training data and used to predict the developability score of the validation set.

## Results

### Knowledge-driven approach for developability prediction

The overall aim of this work was to guide early-stage drug development with algorithms that allow fast selection of a broader set of lead candidates, supplementing and ultimately replacing the high-throughput analytical measurements needed to assess the developability of novel small protein drug candidates such as Anticalin proteins. To do so, it is desirable to define a score that holistically captures the drug-like properties of a protein drug candidate.

A developability target profile was established, beginning with the identification of key developability attributes. Subsequently, critical variables were identified as surrogates for the attributes and assigned accordingly (Figure 1).

Computation of a developability score for each candidate enables a quantitative assessment of that candidate to meet the developability target profile. This developability score was calculated considering multiple analytical measurements, since none of the critical variables alone can provide information about high or low developability risk (Supplementary figure 1). The values obtained for a candidate’s individual critical variables were transformed and normalized whenever applicable, considering the desired target (minimum or maximum) as mentioned in the materials and methods section. The final developability score was calculated as the average of all 17 critical variables.

Figure 2-A shows the scores of the two first principal components after performing a principal component analysis on the critical variables. PCA allows to visualize the dimensionality reduction of the dataset. The information of the 17 critical variables used to calculate the developability score for each construct can be reduced to 2 variables only (the principal components in each axis) for easier visualization, covering 33% of the variability in the original data. To understand how the developability score fits in this reduced dataset, the constructs were colored according to their developability score. To appreciate which variables are influencing the construct’s distribution in the scores plot for these 33% of variance, the loadings plot can be analyzed (Figure 2-B).

**Figure 2.**
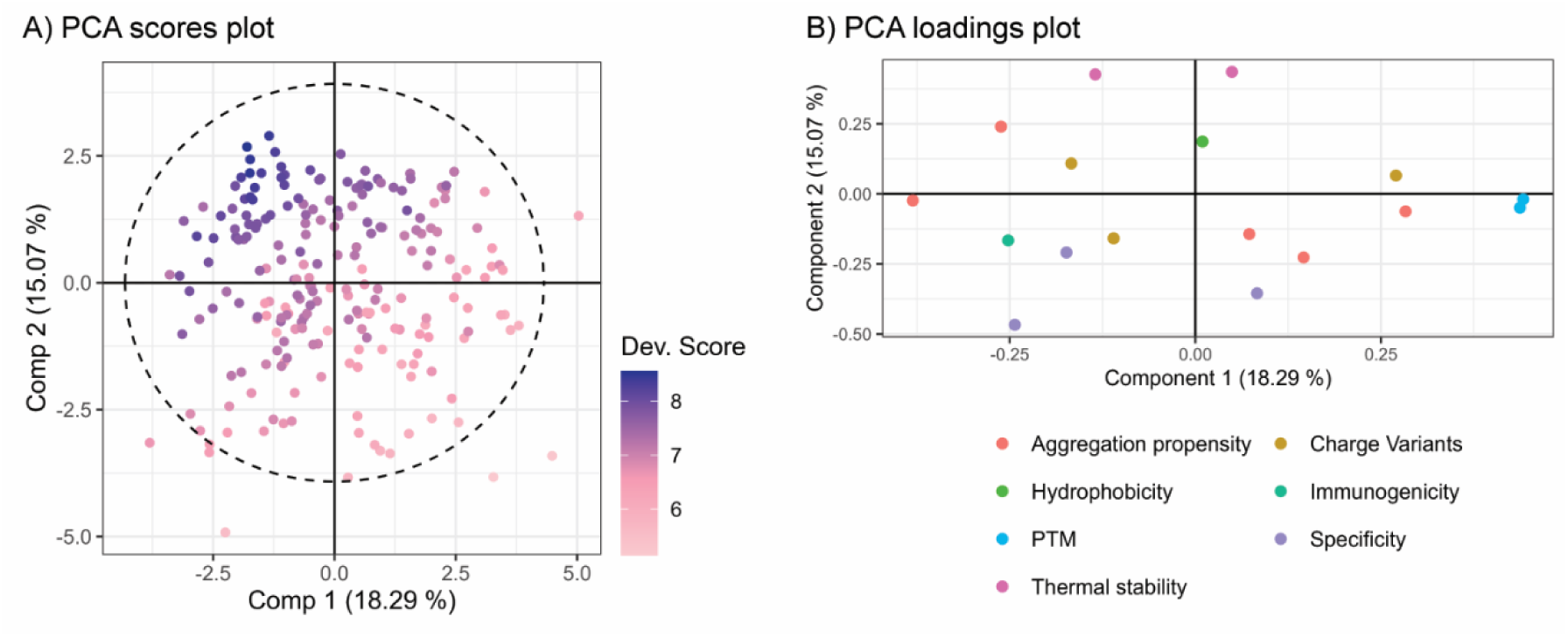
Principal component analysis of the critical variable data used for score calculation. A) Scores plot showing the two first principal components. Constructs are colored by the developability score. B) Loadings plot for the two first principal components. The variables are colored according to the developability attribute they were assigned to.

The loadings plot presents information regarding variable collinearity in the multivariate space. Apart from the two PTM-related variables (blue), the remaining variables in this representation, comprising 33% of the dataset variability, are scattered across the design space with low correlation between each other. This means the choice of critical variables encompasses the full design space without having redundancy in the chosen variables.

### *In silico*-based developability score prediction

After definition of the developability scoring approach and calculation of each construct’s developability score, data-driven models were developed to predict the developability score from *in silico* predictors only (Figure 3). These *in silico* predictors were amino acid sequence-based predictors obtained from the publicly available R package protr (Xiao et al., 2015) and homology model-based predictors calculated using the Molecular Operating Environment (MOE) software (MOE, 2022). Both protr and MOE rely on calculation of protein properties from the amino acid sequence. These properties can then be correlated with developability attributes, such as protein stability or aggregation propensity, attributes that were considered when developing the developability scoring approach.

**Figure 3.**
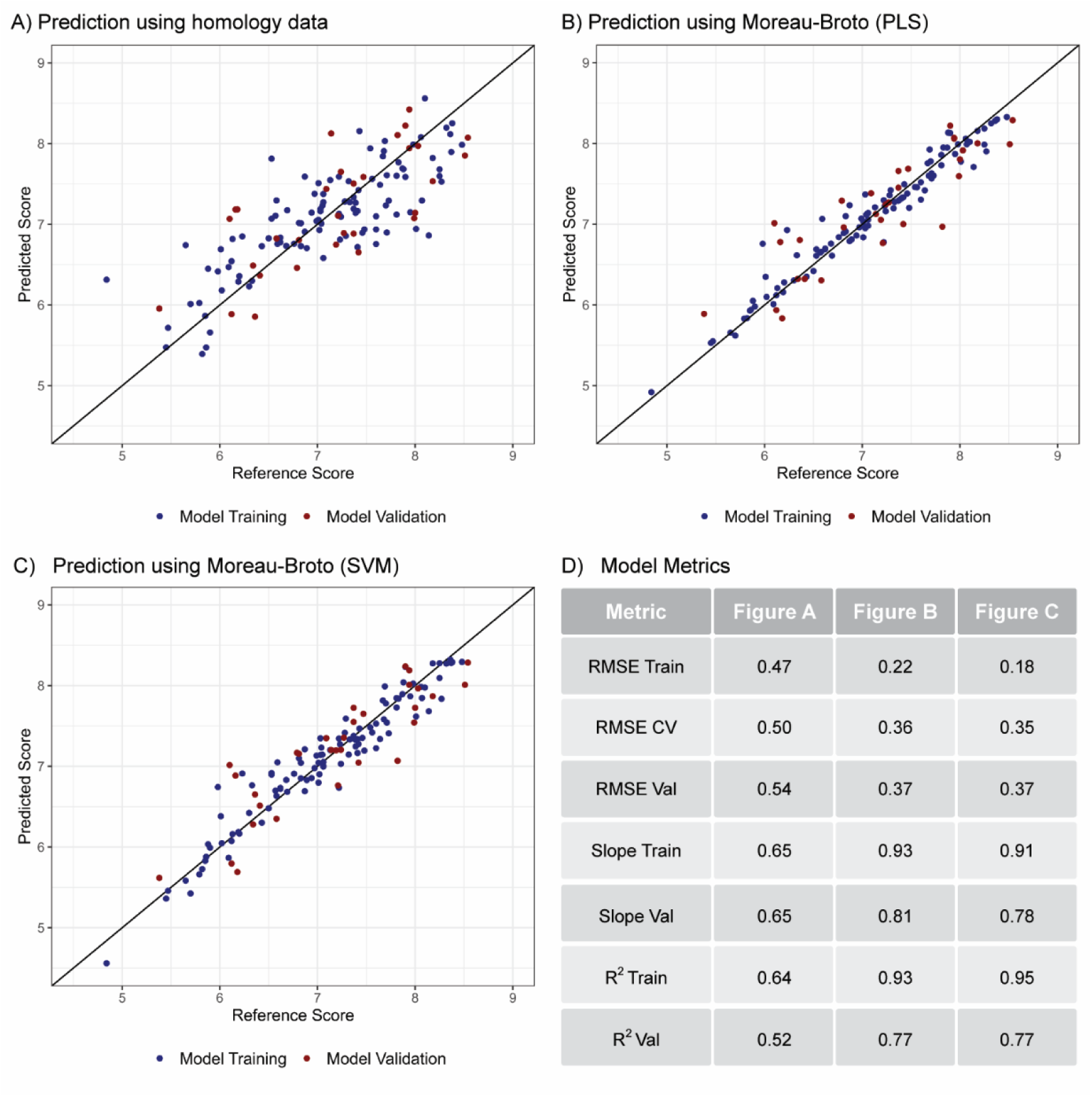
Predicted and calculated developability scores for the training (blue) and validation (red) datasets. A) PLS model prediction of developability score based on MOE homology data, using 2 latent variables. B) PLS model prediction of developability score based on protr sequence-based predictors (specifically, the Normalized Moreau-Broto descriptors), using 6 latent variables. C) SVM model prediction of developability score based on Normalized Moreau-Broto descriptors, based on a first-degree polynomial, with 1 as regularization parameter and scaling parameter of 0.01. D) Model metrics for Figure 3 A-C. RMSE = Root Mean Squared Error; CV = Cross-validation; Val = Validation. Training set size = 105 and validation set size = 31 for all models.

The tested models included partial least squares (PLS) regression and support vector machine (SVM) regression using nonlinear kernels, thus encompassing linear and non-linear modeling approaches, respectively. For the protr descriptors, the approach relies on testing not only each one of the available protr families of descriptors, but also using all protr descriptors simultaneously.

The best prediction results obtained with both model approaches are represented in Figure 3. The normalized Moreau-Broto family of descriptors was the protr family for which better prediction models were obtained. These descriptors are based on the distribution and autocorrelation of eight amino acid properties along the primary sequence (Moreau & Broto, 1980; Xiao et al., 2015).

The models obtained for prediction of the developability score of the validation constructs yielded a validation RMSE of 0.54 for the MOE-based PLS model, and 0.37 for normalized Moreau-Broto PLS and SVM models.

### Leave one target out developability score prediction

A given drug target (i.e., the protein against which a selection campaign is performed) defines the sequence space for functional drug candidates. To assess the model’s ability to predict Anticalin proteins for targets not used in the training process, and thus to simulate the model behavior in production, the leave one target out (LOTO) validation strategy was used. This approach consists in training the model with all available targets except one, and then use that model to predict the left-out target. Since each target has specific signatures for the protr predictors, removing one target leads to an underrepresentation of the sequence space significant for the left-out target, disregarding the predictors more important for that target. To address this issue and reduce the possibility of overfitting, a variable elimination step was introduced, using a recursive feature elimination algorithm. The aim was to select a subset of the 240 Moreau-Broto variables that better describe the training data for all targets, without having too many variables that would lead to overfitting when performing LOTO validation. Variable selection was performed for PLS and SVM, with PLS models yielding better results. Target 7 was left out from the variable selection process, to have an external validation target to assess model predictive ability. The LOTO predictions for the selected PLS model with 40 variables are shown in Figure 4.

**Figure 4.**
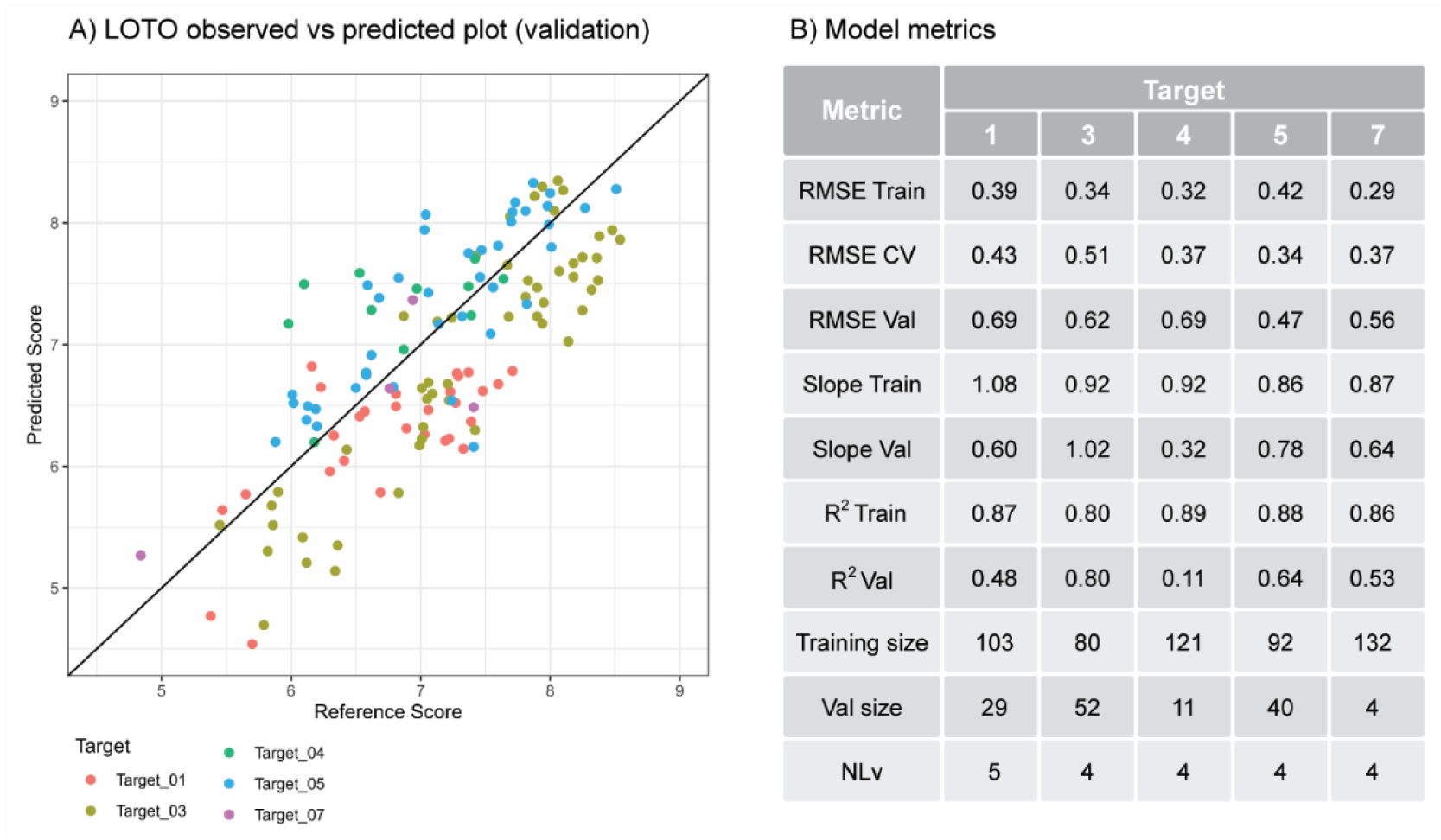
A) Observed and predicted developability scores for the PLS models using RFE- selected predictors from the Moreau-Broto Normalized descriptors. The dots represent the predictions for the target left out. For instance, the target 1 predictions represented in the plot were obtained after training the model with the remaining data. B) Model metrics for Figure 4-A. RMSE = Root Mean Squared Error; CV = Cross-validation; Val = Validation; NLv = number of latent variables.

Since a low RMSE was obtained for the validation set, the approach was tested using the full set of protr descriptors (1520 variables), rather than only the normalized Moreau-Broto, and performing the RFE for each target individually. Running the RFE algorithm only with all the other targets simulates what would occur in the production environment. This resulted in a different number of variables for each target left out. For all targets, PLS and SVM models were tested, with PLS yielding the best results for the target left out. PLS predictions can be observed in Figure 5.

**Figure 5.**
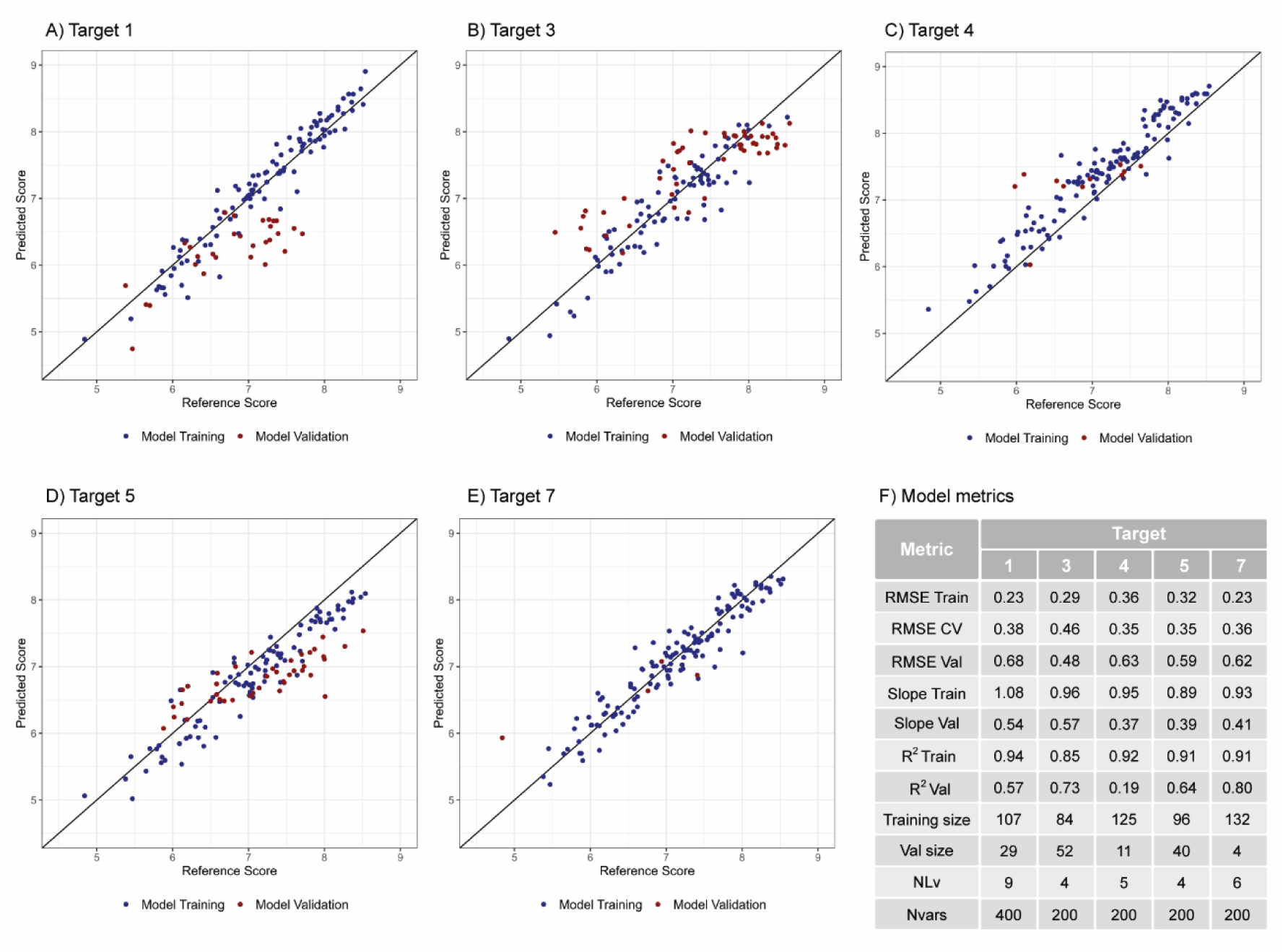
A) to E) Observed and predicted developability scores for the PLS models using RFE-selected predictors from all protr descriptors. For each plot, the training (blue) and validation (red) datasets are shown. The validation set is the target left out during model training, indicated by the plot title. F) Model metrics for Figure 5-A to E. RMSE = Root Mean Squared Error; CV = Cross-validation; Val = Validation; NLv = number of latent variables; Nvars = Number of protr descriptors included in the model.

For each target, the best models have a low number of variables compared to the initial 1520 protr descriptors, with the majority having 200 variables, indicating this variable selection step is important to reduce the available predictors to the ones capturing relevant information from the dataset. Predictions for target 7 are comparable to the remaining targets, indicating the ability of the model to predict an external target not used for variable selection. The results obtained with this approach confirm its validity, and that the models can be optimized if a variable selection step is performed to prevent overfitting.

Another approach tested was the use of hybrid models. These were obtained by considering data from DLS/SLS, *in silico* variables and the protr descriptors. DLS/SLS was chosen since it is a method which can be performed in a plate-based high-throughput format with little evaluation effort, contributing with four critical variables to two developability attributes (Figure 1). A validation RMSE of 0.23 was obtained, lower than the validation RMSE of 0.37 obtained with the Moreau-Broto descriptors only. Since this model was based on 15 principal components summarizing the full 1520 protr descriptors, it is possible the reduction of the noise present in the data culminated in a lower prediction error. However, using the principal components rather than the original variables itself adds an extra layer of complexity to the models. The use of hybrid models (based on *in silico* predictions and selected analytical data) will be further explored in the future.

## Discussion

Prediction of individual developability attributes for standard IgG frameworks is facilitated by the existence of large publicly available datasets that connect analytical characterization with antibody amino acid sequence. However, these data are heterogeneous in nature, originating from different experimental setups including different analytical methods and different production conditions (Khetan et al., 2022), which hampers their usage in the development of predictive models for a targeted developability profile. To develop predictive developability models, it is necessary to start with analytical data obtained from well-defined, comparable experimental setups (Hebditch & Warwicker, 2019).

Relying on a well-defined dataset from the analytical characterization of Anticalin proteins and *in silico*-derived properties, a knowledge-driven approach for developability estimation was established. The chosen methodology is inherently adaptable, allowing its use for proteins based on different small protein scaffolds (e.g., VHH). Moreover, the approach can be further adapted and optimized by adjusting the developability target profile and/or by tweaking the weights of each developability attribute or the individual critical variables for calculating the developability score.

We have calculated this holistic developability score for the set of drug candidates used in this study as it offers a simplified yet pragmatic approach to capture a complex system of various biophysical and physicochemical properties. Certainly, an accumulated developability score does not give detailed information on each individual developability attribute. However, the intention of this approach is to guide early stages of discovery, where the task is to filter for candidates with the highest scores from hundreds or thousands of sequences identified from a screening campaign or in silico engineering approaches. Thus, exploring the specific factors contributing to medium or low performing candidates is of minor concern at this stage. It is acknowledged that later in-depth developability assessments of a small number of final candidates require an individual evaluation of the different developability attributes.

The sequence-based models developed in this work predict the Anticalin protein developability score with RMSE ranging from 0.37 to 0.69. Considering the entire range of developability scores from approximately 5 to 8.5, this corresponds to a percentage error of ∼10 to 20 %. The finding that amino acid sequence can be predictive of developability for antibodies was already explored by other authors (Chen et al., 2020; Hebditch & Warwicker, 2019). Chen and co-authors used embedding classification models based on the antibody sequence to predict the developability index (Lauer et al., 2012), which accounts mainly for aggregation propensity. However, the authors mention their models were overfitting. In this work, sequence-based calculated properties were correlated with a developability score which considers seven key developability attributes to be assessed in early development. Overfitting was prevented by building robust models using the recursive feature elimination algorithm and leave one target out validation was conducted to confirm models were not overfitted. Hebditch and Warwicker relied on 35 sequence-based features to build models trained on the Jain et al. dataset (Jain et al., 2017) and obtain predictions for 12 analytically measured properties related to developability, such as differential scanning fluorimetry or expression titer (Hebditch & Warwicker, 2019). These models can be used for relative ranking of the candidates. The authors reported R^2^ values up to 0.391 in their models. Herein, the models presented in this work (Figure 3) have R^2^ values between 0.52 and 0.77. Since the models were validated using similar approaches and the dataset size is very comparable, two possible explanations for the overall better models obtained here are the differences between Anticalin and antibody size and sequence lengths, and the number of sequence-based predictors used for modeling (1520 vs 35).

When comparing protr and MOE-based predictions, a better prediction was obtained with the former. This might be due to the number of descriptors obtained in each approach, which is significantly higher for protr. Moreover, the fact that MOE descriptors are calculated based on homology models and that such models are least accurate for the flexible loop regions which are the main discriminators between the individual Anticalin proteins, may hamper the use of those descriptors. In the future, MOE-derived data will also be explored with the use of feature engineering techniques to identify the most relevant descriptors for the developability score. Despite the better results obtained, a disadvantage of using *in silico* descriptors such as protr, is that the impact of the formulation in the descriptors’ results is not considered (Hebditch & Warwicker, 2019; Lauer et al., 2012; Narayanan, Dingfelder, Butté, et al., 2021). This impact is minimized if the developability score establishment and the developability score prediction are performed in the same developability stage, with the same analytical methods and formulation conditions, as performed in this work. Still, it might be worth exploring both protr and MOE descriptors and select the ones more tailored to the final pharmaceutical formulation.

In this work, in particular for the leave one out validation, PLS models were chosen as the final models. Bailly et al. (Bailly et al., 2020) discussed the usefulness of the predictive potential of PLS models to capture relevant information from multiple predictors and increase the model’s robustness. The fact that PLS models’ performance was better than the SVMs’ indicates the interactions in the dataset are mostly linear. SVM presented a tendency to fit the training data better than PLS models, but afterwards the validation error was higher.

Importantly, the models yield acceptable predictions when the construct to be predicted has a score in the range used for model calibration. But when the score falls outside of this range, as is the case for one of target 7 constructs (Figure 5-E), the score prediction is impacted, demonstrating the importance of having a model calibration range spanning through low and high scores. It can also be observed throughout the models tested in this work (Figure 3-B and C) that in general there is a difference in the ability of the models to predict scores below 6.5 and for scores above that threshold, with latter predictions being more accurate. This likely stems from having more training data available with a developability score above 6.5 (approximately 79% of the training data). When new data is available and constructs with lower developability scores can be analyzed and added to the model, it is expected that the model prediction ability for the lower scores will also improve.

Regarding the leave one out validation strategy, it can be concluded that the quality of LOTO predictions is target-dependent, with targets 3 and 5 being the ones for which better predictions are obtained (Figure 5 B, D and F). It is also interesting to observe that target 3 is consistently grouped away from the remaining targets in the PCA plots, with a separated cluster on PC1 (Supplementary figure 2), and it is the one with higher representation (38%) in the training data. These factors likely contribute to a higher influence of target 3’s protr descriptors on the model, which leads to overfit and hindering other targets’ predictions in the LOTO models using normalized Moreau-Broto (results not shown). Due to the reduced dataset size, common to biological datasets, removing target 3 from the analysis further worsened predictions, meaning the data added by including proteins binding this target are providing important information to the model.

By reducing the number of variables to include in the models using the RFE algorithm, focusing on the parameters capturing information from all targets, model predictions for the target left out could be improved (Figure 4 and Figure 5). Since variable selection for normalized Moreau-Broto was performed with the presence of all targets except target 7, predictions for this target represent the situation verified in production, when a completely new target is produced and its developability score must be assessed. The results obtained are good for production purposes and comparable to the remaining targets.

As shown in Figure 5, the RFE strategy can be applied to optimize the model to a specific target. This demonstrates that *in silico* descriptors can be used for accurate prediction of the developability of early-stage candidates. As Anticalin proteins against more targets are generated, characterized and added to the training set they will increase the variability of the dataset. The variables selected will be able to capture the main characteristics of the *in silico* data that better correlate with the developability attributes, yielding robust models able to generalize and predict accurate developability scores for new targets. Analyzing the model calibration and prediction metrics as each new target is added to the model in production will allow to understand when the variables selected by RFE are representative of the global Anticalin variety and performing RFE for each new target is no longer necessary.

With this work, the validity of the explored strategy is confirmed, since the models obtained have a low RMSE, and are robust and reliable enough to be utilized in a live production system. A future pipeline will generate more data, increasing the number of different targets present in the calibration set, to increase model’s exposure to different Anticalin protein variants and enhance model performance.

The developability scoring approach devised in this work can possibly support the field of biopharmaceutical informatics, enabling a higher probability of success in developing targeted protein therapeutics. Further investigation of hybrid models and identification of the most critical analytical assays allows to streamline analytical characterization, by reducing the number of analytical assays or methods and consequently saving time and resources. Additionally, a complete *in silico* sequence-based prediction, independent of analytical assays, allows to rapidly screen candidates to select the ones to move to a next developability stage, where analytical characterization can take place, once again saving valuable time and resources.

## Conclusions and future steps

In this work, we have devised a holistic approach to calculate the developability score of Anticalin proteins, allowing for an unbiased ranking of drug candidates and an accelerated workflow in early-stage drug development. This flexible approach allows weighing of individual attributes to adapt the approach to other protein scaffolds or to incorporate emerging pipeline knowledge on a given modality or route of administration.

By calibrating PLS and SVM models on this developability score and on a set of *in silico* descriptors, the developability score for new constructs was predicted with good accuracy. As new data is generated, it will be added to the models in production to further improve model robustness and increase model’s predictive ability. Having established the workflow for developability score prediction and studied the ranges obtained, the efforts are now focused on defining the limit separating a promising candidate from a bad one (Bailly et al., 2020). Once this limit is specified, the models can be adapted to return the probability of a candidate to proceed to the next developability phase.

To explore the different factors involved in the developability score calculation and *in silico* model training and prediction, an interactive app was developed. The app was designed to allow the user to test different constructs, targets and models, including the possibility to explore analytical and *in silico* variables. Additionally, the possibility to calculate a developability score for each developability attribute was considered, by averaging the critical variables of each attribute. This way, model-based predictions of each developability attribute can be explored. Different modeling algorithms were included to assess the various prediction capabilities for new data, allowing the user to test for instance PLS and SVM models and choose the best one. Model lifecycle management can also be performed interactively, allowing the user to recalculate the new models as more training data is obtained. Finally, this tool can be used to generate previously unidentified variants during the selection process and generate new libraries, effectively excluding low scoring Anticalin proteins from the beginning of the candidate selection process.

## Acknowledgements

We would like to thank all colleagues from the Pieris Pharmaceuticals GmbH Protein Analytics, Pre-Clinical and Technical Development teams who supported this project by generating protein materials and the analytical data. Especially we thank Stefan Grüner for calculating the MOE descriptors and Epibase immunogenicity scores, Jéssica Pestana and Susana Lousa for preparation of the graphical abstract and helping with figure preparation and Rui Silva, Rui Almeida and Ângela Martinho for critical review of the manuscript.

## Author Contributions

Daniel: Conceptualization, Methodology, Software, Validation, Formal Analysis, Data Curation, Writing – Original Draft, Writing – Review & Editing

Jan-Peter: Conceptualization, Software, Data Curation, Writing – Original Draft

Karin: Conceptualization, Investigation, Data Curation, Writing – Review & Editing, Supervision

Maria: Methodology, Software, Validation, Data Curation, Writing – Review & Editing

Timo: Conceptualization, Writing – Review & Editing

Sofia: Conceptualization, Methodology, Software, Validation, Formal Analysis, Data Curation, Writing – Review & Editing

Raman: Software, Writing - Review & Editing

Sandra: Conceptualization, Methodology, Software, Validation, Formal Analysis, Resources, Data Curation, Writing – Original Draft, Writing – Review & Editing, Supervision

Hitto: Conceptualization, Resources, Writing – Review & Editing, Supervision

