## Supplementary Data for "Holistic in silico developability assessment of novel classes of small proteins using publicly available sequence-based predictors"

### Supplementary tables

*Supplementary Table 1 - Amino acid sequence motifs for each post-translational modification considered. PROSITE patterns are used* (Sigrist et al., 2012)*. Amino acid abbreviations: A – Alanine; D – Aspartate; G – Glycine; H – Histidine; M – Methionine; N – Asparagine; P – Proline; R - Arginine; S – Serine; T – Threonine; W – Tryptophan.*

| Post-translational modification motifs | Amino-acid combinations |
| --- | --- |
| Deamidation | N[GSTNH] |
| Isomerization | D[GSTDH] |
| Fragmentation | DP |
| Integrin-binding | RGD |
| Oxidation | [WM] |
| N-glycosylation | N{P}[ST] |
| O-glycosylation | [PSTA][PSTA][ST][PSTA][PSTA] |

### Supplementary figures

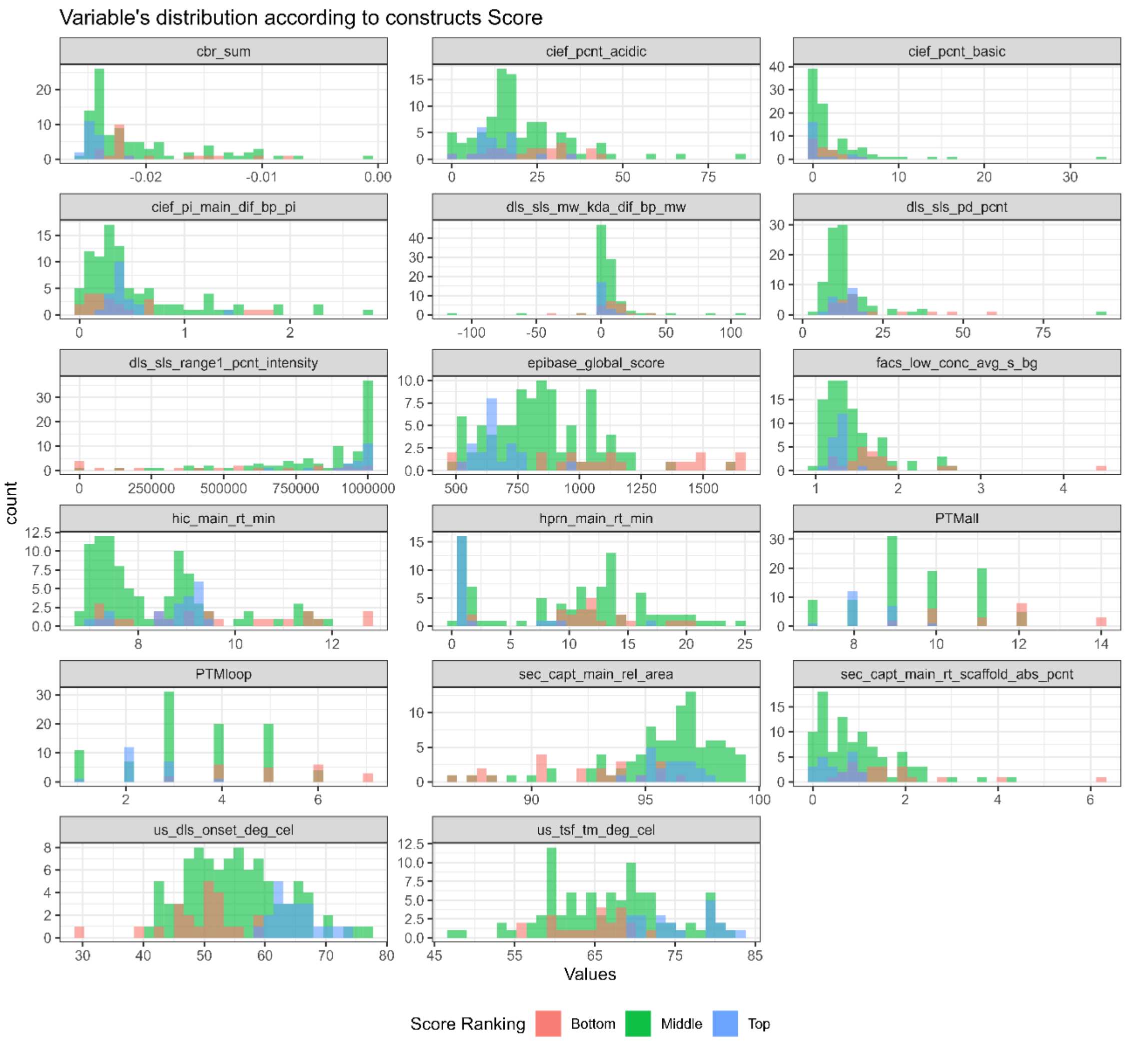


*Supplementary figure 1 – Distribution of the values obtained for each of the 17 critical variables depending on whether the Anticalin candidate's developability score belonged to the top (blue), middle (green) or bottom (red) range.*

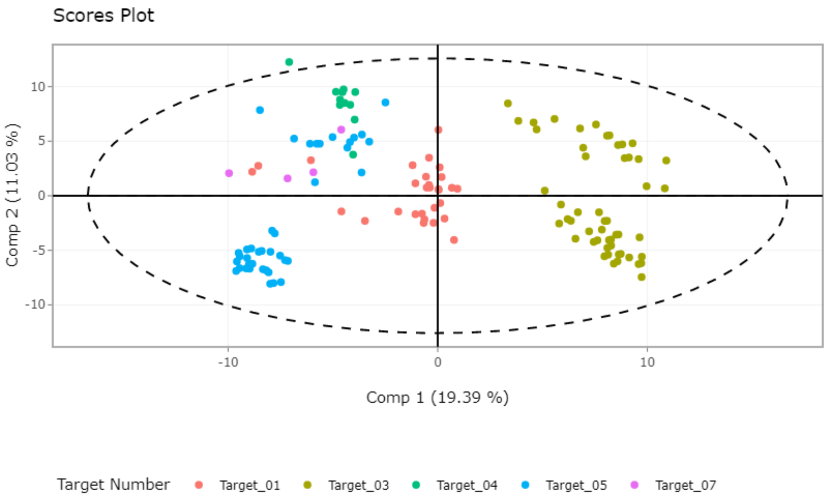


*Supplementary figure 2 - PCA scores plot of the Moreau-Broto descriptors colored by each of the five targets.*
